# Standard touchscreen test procedures shift circadian rhythms in mice: a modified protocol to minimize circadian confounds

**DOI:** 10.64898/2026.10.27.762380

**Authors:** Alice I Ayres, Taha Yildirim, Samantha Saw, Mateusz Michalik, Lea Paty, Mayuko Arai, Robert Gibson, Ralph E Mistlberger, Brianne A Kent

**Affiliations:** Department of Psychology, Simon Fraser University; Institute for Neuroscience and Neurotechnology, Simon Fraser University; École supérieure de physique et de chimie industrielles de la ville de Paris

**Keywords:** Circadian rhythms, touchscreen, pattern separation, TUNL, cognitive testing, rodents, photic entrainment

## Abstract

Touchscreen-based cognitive testing for rodents offers considerable experimental benefits, including a semi-automated battery approach and the use of appetitive rather than aversive reinforcement. However, food-reward and touchscreen illumination introduce potential confounds, as both food and light can shift, entrain, or disrupt circadian rhythms, which in turn can affect learning and memory. The potential for the test apparatus to impact endogenous circadian rhythms is particularly problematic when probing relationships between sleep, circadian disruption, and cognitive performance. After observing phase shifting of circadian rhythms in nocturnal mice tested using standard touchscreen test procedures, which include testing during the active period (early lights-off), we evaluated the relative contributions of the illuminated screen and the appetitive food reward. Our results indicate that light emitted by the touchscreen chambers is responsible for the circadian phase shifting effects, with standard screen conditions inducing a ∼3.5 h phase delay in daily activity rhythms when training is conducted early in the dark period for 7 consecutive days. We then modified lighting in the test environment and the home cage to identify parameters that would minimize circadian shifting without impairing visual discrimination or performance on a widely used cognitive test (Trial-Unique Nonmatching-to-Location, TUNL). From the 10 conditions tested, we offer specific recommendations for modifying touchscreen test protocols to minimize circadian confounds, without impacting task acquisition or performance.

**Highlights:** * Daily touchscreen testing shifts circadian rhythms in mice

* Photic, rather than food-related, cues underlie the phase shifting effect

* Modifying touchscreen settings can minimize the effects on circadian rhythms

* Protocol modifications that reduce phase shifting do not compromise TUNL performance

**Data Statement:** Data will be available upon publication through the SFU Institute for Neuroscience and Neurotechnology data portal.

## 1.0 Introduction

Touchscreen-based procedures to test cognitive processes in lab animals have significant experimental advantages over other behavioural measures of cognition (e.g., maze-based tasks), including a semi-automated battery approach, high reproducibility across experiments, precise stimulus control, sensitivity to subtle cognitive differences, appetitive rather than aversive reinforcement, compatibility with neurotechnology such as optogenetics, and strong potential for insights to translate to humans [1]. Importantly, the touchscreen operant platform permits assessment of targeted cognitive domains comparable to the test battery used in humans (Cambridge Neuropsychological Test Automated Battery, CANTAB) [2].

Cognitive assessments in humans are typically conducted during the day, unless there is special interest in effects of time of day or sleep-targeting manipulations. Nocturnal mice are the most used animal models [3], and many labs maintain mice under inverted light-dark (LD) cycles so that behavioural tests can be conducted during the daily wake phase of both the test subjects and the investigators. Touchscreen cognitive tests employ light to deliver visual stimuli and restricted feeding schedules to support food-reinforced training and performance. The daily, timed exposure to light and food stimuli raise methodological issues related to how test procedures affect circadian rhythms, which to date have been largely overlooked.

The endogenous circadian clocks that regulate behaviour and physiology in mammals are entrained to local time by sensitivity to photic and non-photic time-cues (zeitgebers) [4–6]. The response to these time cues varies with time of day and stimulus parameters. Light, the dominant zeitgeber, has relatively little effect during the day (sleep phase in nocturnal rodents), but shifts circadian rhythms to an earlier phase (cycle delayed) when exposure occurs late in the day and early in the night, and to a later phase (cycle advanced) when exposure occurs late in the night and early in the day. Non-photic stimuli, among which food intake is most important, can also shift circadian clocks in many organs and tissues, and together with light, establish an internal circadian program that aligns rhythms to the solar day at a characteristic day- or night- active phase. Circadian rhythms in nocturnal rodents are exquisitely sensitive to light during their biological night, and even brief exposure to a few photopic lux can induce phase shifts [4, 5]. Whether light exposure required for touchscreen testing during the dark phase might shift rhythms has not been assessed to our knowledge, despite evidence that cognitive processes are sensitive to time of day, and impaired by disruptions of circadian rhythms and sleep (reviewed by [7–9]).

Here, we investigated potential effects of daily touchscreen cognitive testing on circadian activity rhythms in mice. This inquiry followed our observation that Trial-Unique Nonmatching-to-Location (TUNL) testing early in the dark phase shifted circadian activity rhythms in mice that were entrained to a standard light-dark (LD) cycle and impeded re-entrainment to a 6-h delay of the LD cycle. Standard touchscreen conditions expose mice to polychromatic white light of up to ∼21 photopic lux (with each stimulus square on the screen emitting an additional ∼7.5 lux), which are intensities sufficient to phase-shift C57Bl6 mice [10]. Daily touchscreen testing also requires a palatable food reward (typically a strawberry milk drink), which can act as a zeitgeber for the circadian system [11, 12], and mild daily caloric restriction to ensure motivation to work for food reward. We therefore evaluated the independent contributions of the touchscreen light and palatable food reward to the phase shifting effect.

We confirmed that exposure to the touchscreen chamber (including photic and non-photic task stimuli) for an hour beginning 2.5 h after lights-off can markedly alter the phase of entrainment to the LD cycle, which effectively changes the (subjective) circadian time of the test sessions. We next established that touchscreen-emitted light was the primary stimulus responsible for the phase shifting effects. We then explored three approaches to minimize this effect, by (1) reducing screen or stimulus brightness, while maintaining sufficient illumination to perform visual discrimination tasks, (2) red-shifting the emitted light, to exploit the relative insensitivity of the rodent circadian system to long wavelengths [10], and (3) increasing home cage light intensity, which may enhance circadian clock amplitude and thereby decrease the magnitude of light-induced phase-shifts [13, 14]. Finally, we assessed the impact of chamber modifications on TUNL performance and provide practical recommendations for touchscreen- based testing of nocturnal rodents during their active phase to minimize circadian rhythm impacts.

## 2.0 Methods

### 2.1 Animals

C57BL/6 mice were obtained from Charles River Laboratories (Quebec, Canada) and adapted to the local light:dark cycle (LD 12:12) in group cages for several weeks. The mice were habituated to handling before use in experimental procedures. Experiment 1 used 22 mice (12 females; 22 weeks old), Experiment 2 used 20 mice (10 females; 14 weeks old), Experiment 3 used 8 mice (4 females; 9 weeks old), and Experiment 4 used 20 mice (10 females; 7 weeks old). All procedures complied with Canadian Council on Animal Care standards (Simon Fraser University ethics protocol 1325P-21).

### 2.2 Housing and activity data collection

Mice were housed in standard polycarbonate cages (20.5 x 14 x 36.5 cm) with stainless steel wire-bar tops, lined with enriched cob substrate, and containing tinted acrylic igloos (Igloo Fast-Trac, BioServ, USA; 11 cm diameter, 5.7 cm tall) with shredded paper for nesting. Cages were kept in light-proof and sound-attenuating cabinets. An inverted 12:12 LD cycle was implemented to facilitate testing during the dark period. Light (∼70 lux) was provided by white LEDs mounted over each cage. Temperature was maintained at 21 ± 1 °C with controlled humidity. Females were housed in pairs for experiment one and singly for experiments 2-4.

Males were housed individually. Locomotor activity was detected using passive infrared motion sensors mounted above each cage and monitored continuously with the ClockLab data acquisition and analysis system (Actimetrics, IL, USA). Water was available ad libitum in the home cage and food was provided each day 7 h after lights-off (Zeitgeber Time (ZT) 19, where lights-off is ZT12 by convention), in an amount sufficient to maintain body weight at 90–100% of free feeding weight during touchscreen training and testing. Mice were weighed daily during food restriction. On no-test days, and during constant darkness (DD) mice were left undisturbed in home cages until feeding time (experiment one) or with food *ad lib* (experiments 2-4). To minimize circadian disruption, health checks and biweekly cage changes were performed briefly under red light during their active period. When mice were transported to the testing apparatus, they were transferred through the facility in light-proof boxes and the housing and testing rooms were dimly lit with red light (∼8 lux and 2.5 lux, respectively).

### 2.3 Touchscreen Apparatus

Four 2nd-generation Bussey-Saksida Mouse Touchscreen Chambers were used (Campden Instruments Ltd., Loughborough, UK; model 80604A/614A). Each animal was assigned to a single chamber for the duration of the experiment; chambers were cleaned between animals to remove olfactory cues. Each touchscreen was fitted with a mask exposing a single row of six response windows (each window was 5 cm x 3.5 cm). Task schedules were designed, run, and recorded using Whisker Server and ABET II software (Lafayette Instruments), adapted in-house from TUNLv2 schedules [15]. The food reward provided was a strawberry flavoured nutritional supplement shake (Ensure Regular, Abbott, Saint-Laurent, Canada), which we refer to as “milkshake.”

### 2.4 Touchscreen Pretraining

Pretraining was conducted as previously described [16, 17]. Briefly, pretraining progressed through (1) introduction to food reward in home cage for 2 days, (2) habituation to the testing chamber for 10 min, (3) habituation to the milkshake in the testing chamber for 20 min one day and 40 min the next, (4) Initial Touch (60 min, 30 trials), where mice were trained to associate a stimulus (illuminated white square) in one of six response locations with food reward delivered at the magazine, (5) Must Touch (60 min, 30 trials), where mice were trained to touch the stimulus directly to obtain food reward, (6) Must Initiate (60 min, 30 trials), where mice were trained to initiate each trial by breaking the magazine IR beam, (7) Punish Incorrect (60 min, 48 trials), where mice were trained that only touches to the illuminated stimulus were rewarded.

### 2.5 Cued TUNL

Once mice successfully completed pretraining, they were trained on Cued TUNL as previously described [15]. Briefly, the sample phase illuminates one of six locations on the touchscreen.

Following a delay, the original sample location (incorrect) is presented with 50% reduced luminance alongside the correct novel location at full brightness, intended to help mice acquire the non-matching rule more efficiently. During a session, mice progress from ‘easy’ trial configurations, with the maximum separation distance (S4), through to the harder separation distances (minimum separation distance = S0) via the within-session ‘stage’ progression system. Mice were trained daily until completing two consecutive sessions of 48 trials at >70% accuracy while reaching separation Stage S1 (S0 Stage was not required).

### 2.6 TUNL Task

Once mice successfully completed Cued TUNL, mice were tested on TUNL as previously described [15]. Briefly, during the sample phase, a stimulus appeared in one of six locations. A correct touch triggered a tone, removed the stimulus, and dispensed the food reward at the magazine. Breaking the magazine beam turned off the magazine light and initiated the choice phase, in which the original sample location and a novel location were both illuminated at equal luminance. Correct identification of the novel location was rewarded with a tone and food reward, followed by a 15-s ITI. Touches to incorrect (blank or sample) locations triggered a correction trial. Trial difficulty was determined by the separation distance between sample and choice locations (S4 = least difficult to S0 = most difficult). The smaller separations increase similarity between locations and are more taxing on pattern separation [18].

## 3 Results

### 3.1 Touchscreen testing alters circadian rhythm entrainment and response to LD shift

Experiment 1 was designed to evaluate the effect of a 6-h delay of the LD cycle on TUNL performance. Mice were assigned to a TUNL-LDshift group (6-h delay) or a TUNL- Control group (no shift) (n = 11/group; females = 6/group, pair-housed). TUNL sessions were scheduled during the first 6 h of the dark period, at one of 6 ZTs (ZT12.08, 13.17, 14.33, 15.5, 16.67 or 17.83). On days with no TUNL sessions, mice were weighed and fed at the usual time (ZT19). To evaluate performance effects of the 6-h LD phase shift, mice were tested for 2 days on S2 stage to establish a performance baseline and then the LD cycle was delayed by 6 h in the TUNL-LDshift group. The LD shift was initiated by extending the dark period by 6 h for one cycle so that lights-off began 6 h later (phase delay shift, comparable to translocating 6 times zones west). Mice were tested on the S1 stage for 9 of the first 10 days following the LD shift and then maintained in DD for 2 consecutive days without testing, to assess re-entrainment to the shifted LD cycle.

Visual inspection of actograms indicated that for some mice the touchscreen test procedure can substantially alter the timing of nocturnal activity and re-entrainment of circadian rhythms to a shifted LD cycle (Fig. 1A-D). To quantify and interpret these effects, the onset of nocturnal activity was assessed on days without testing, either in LD or DD (periods of one or more days). As an effect of touchscreen testing on circadian rhythms was not anticipated, a separate group of undisturbed control mice naïve to the touchscreen protocol were not included. Therefore, archival data were used for comparison. These data were obtained from two groups of *ad-libitum* fed young adult male C57BL/6 mice previously recorded in the same cages and naïve to touchscreen testing. One of these groups was recorded in LD and 10 days of DD (AL-DD group), and the other in LD before and after a 6-h LD delay shift (AL-shift group).

**Figure 1.**
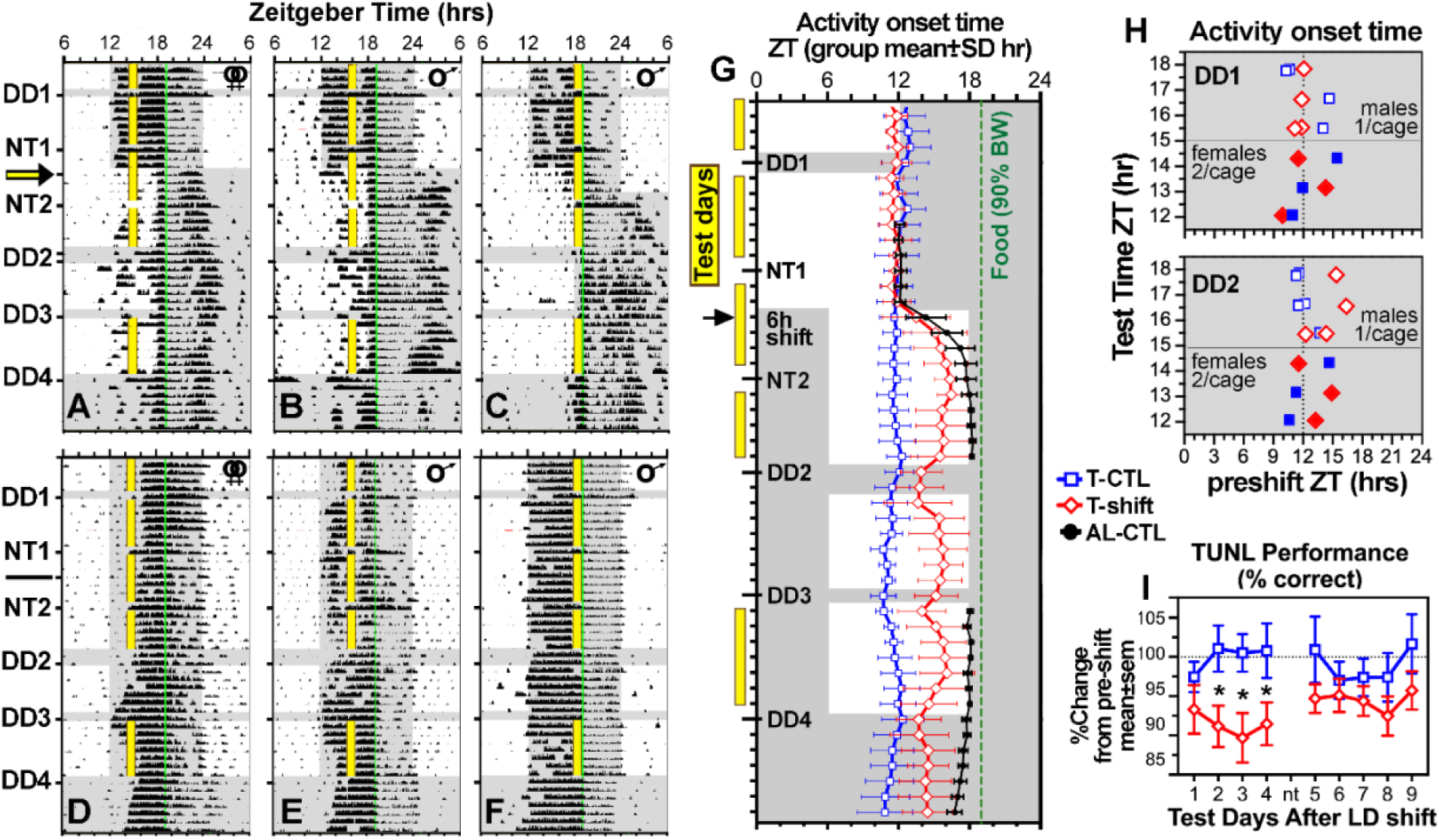
Circadian activity rhythms and TUNL performance of mice in Experiment 1. **(A-F)** Activity data plotted in standard ‘actogram’ format from representative pairs of female (**A,D**) and single male mice tested at ZT14.5 **(A, D**), 16.67 **(B, E)** or 17.83 **(C, F).** Five-min bins during which one or more activity counts were registered are indicated by a vertical bar proportional in height to the amount of activity, expressed as a percentile for that day. Each line represents one full 24**-**h day, with consecutive time bins plotted left to right, and consecutive days aligned vertically. Gray shade denotes lights**-**off. Yellow vertical bars denote TUNL test times. A thin green bar denotes the time of day that mice received a daily allotment of chow sufficient to maintain body weights at ∼90% of free-feeding weight. The LD-shift day is indicated by an arrow. **(G)** Group mean (**±**SD) daily activity onset times in the TUNL-LDshift (T-shift, red), TUNL-control (T-CTL, blue) and AL-control (AL-CTL, black) groups, in LD and DD. (**H**) Activity onset times (x-axis) plotted against daily TUNL test times, in TUNL-LDshift (red symbols) and TUNL-control (blue symbols) mice on two days without testing in constant dark (DD), one before the LD shift day (DD1) and one after (DD2). Filled symbols indicate females and open symbols indicate males. **(I)** Group mean (**±**SEM) overall accuracy on TUNL S1 stage sessions of mice in the TUNL-LDshift (red) and TUNL-control (blue) groups during the first 10 days after the LD shift (no test on day 5). Y-axis shows baseline corrected accuracy (percent correct). * denotes significant difference between groups (multiple T-test, *p* < .03). ZT, zeitgeber time; DD, constant darkness with no testing; LD, light-dark; TUNL, Trial-Unique Nonmatching-to-Location; NT, no-test day in LD.

On the last test-free day in LD prior to the LD shift, nocturnal activity onset occurred at ZT12.05 **±** 1.23 h in the TUNL-LDshift mice and at ZT11.78 **±** 0.53 in the TUNL-Control mice). While these group averages are close to that of AL-DD mice (ZT12.10 **±** 0.27), within- group variability was substantially greater, with onsets ranging from ZT9 to ZT15 across mice in the TUNL groups (e.g., Fig. 1D, E), compared to ZT11.5 to 12.7 in the AL-DD group.

After 51 days with near-daily TUNL training, mice in the TUNL-LDshift group were subjected to a 6-h delay shift of the LD cycle. The expected response is a rapid phase delay of the rest-activity cycle, with the nocturnal timing of activity reestablished in about 4 days, as exemplified by AL-shift mice (N=16), all of which were re-entrained by day 5 (Fig. 1G). By contrast, TUNL-LDshift mice showed variable responses, with some mice delaying 6 h (e.g., Fig. 1C) and others shifting only partially (e.g., Fig. 1A, B). On days 10-11 after the shift, the lights remained off for 48 h, and the mice were not disturbed, to assess the phase of entrainment without direct effects of light or the test procedure. Average activity onset time in the TUNL- LDshift mice occurred at ZT13.92 **±** 1.77 h (ZT of the LD cycle before the shift), a delay of only 1.88 h in 10 days. By contrast, activity onset in the AL-shift archival group occurred at ZT18.11 **±** 0.42 h, a full 6-h delay.

Each mouse in the TUNL groups was assigned to one of 6 test times during the first 6 h of the dark period (Fig. 1G). There was no clear association between test time and the phase of activity onset prior to or after the LD shift (Fig. 1 I-L), but due to small sample sizes and unequal representation of males (single housed) and females (pair housed) at the different test times, no further analyses were attempted.

To evaluate performance effects of the 6-h LD shift, both groups of TUNL mice were tested on S2 stage for 2 days to establish a performance baseline, and then on S1 stage on 9 of the 10 days after the LD shift. As expected, both groups scored lower on the more difficult S1 stage, but the decrement was significantly greater in the LD-shift group. Repeated measures ANOVA confirmed a significant main effect of group (F_1,20_ = 5.223, p = .033), and no main effect of day (F_5.969,119.4_ = 1.079, p = .3789) or interaction (F_5.969,119.4_ = 0.6728, p = .3789). Group differences were significant on days 2, 3 and 4 after the LD shift (Fig. 1I).

### 3.2 Experiment 2: Dissociating photic and non-photic entraining cues

Experiment 1 revealed an effect of touchscreen testing on entrainment of circadian rhythms to LD, and on re-entrainment following a 6-h LD shift. These effects suggest that photic and/or non-photic stimuli associated with the touchscreen testing procedure can act as circadian time cues that can shift circadian clock phase or modify its response to an LD shift. Experiment 2 examined circadian responses to photic and non-photic stimuli in isolation.

Mice were randomly assigned to four groups (n = 5/group) and each group experienced two or three of the four exposure conditions: (1) *Light condition* in which mice were placed in the touchscreen chamber running a schedule designed to reflect the lighting encountered by a mouse performing at 80% accuracy on the TUNL task (the screen displayed 100%-luminance TUNL task images with intermittent illumination of the magazine and house lights). No milkshake reward or audio cues were present; (2) *Dark condition* in which mice were placed in the touchscreen chamber with the apparatus switched off entirely to remove all light sources, audio cues or milkshake reward; (3) *Milkshake condition* which was identical to *Dark condition*, but with the milkshake pump operational, delivering reward at intervals and volumes comparable to a mouse performing the TUNL task at 80% accuracy; and (4) *Home cage condition* in which mice were left undisturbed in their home cage. Mice experienced 60-min exposures for 7 consecutive days at ZT14.5, followed by DD for 10 days. Exposure conditions were separated by at least 2 weeks, to allow activity rhythms to re-entrain to LD. Activity onset on DD day 1 was used to quantify the phase of entrainment to LD after each experimental condition, and onsets on all 10 days were used to quantify the circadian periodicity (t) of the endogenous activity rhythm. Using the ClockLab offset detection algorithm, we determined the end of each animal’s daily active bout and subtracted each day’s corresponding onset from it to calculate alpha (a; the duration of the active period).

Exposure to the Light condition markedly delayed the onset of nocturnal activity during the week of testing in all 20 mice tested (e.g., Fig. 2A-D). The delayed onset was evident on the first day of DD, when the mice were not disturbed, confirming a delayed phase of entrainment to the LD cycle (Fig. 2E-H). The Dark and Milkshake conditions had little effect by comparison. A mixed effects analysis indicated a significant main effect of condition (F_3, 24_ = 237.7, *p* < .001), no main effect of sex (F_1, 18_ = 0.851, *p* = .368) and no interaction (F_3, 24_ = 2.706, *p* = .068). Post-hoc tests revealed a significant difference between the Light condition and the other 3 conditions in males and females (Tukeys, adjusted *p* < .001), a significant difference between Milkshake and Home cage control conditions in females only (adjusted *p* < .038), and no other differences.

**Figure 2.**
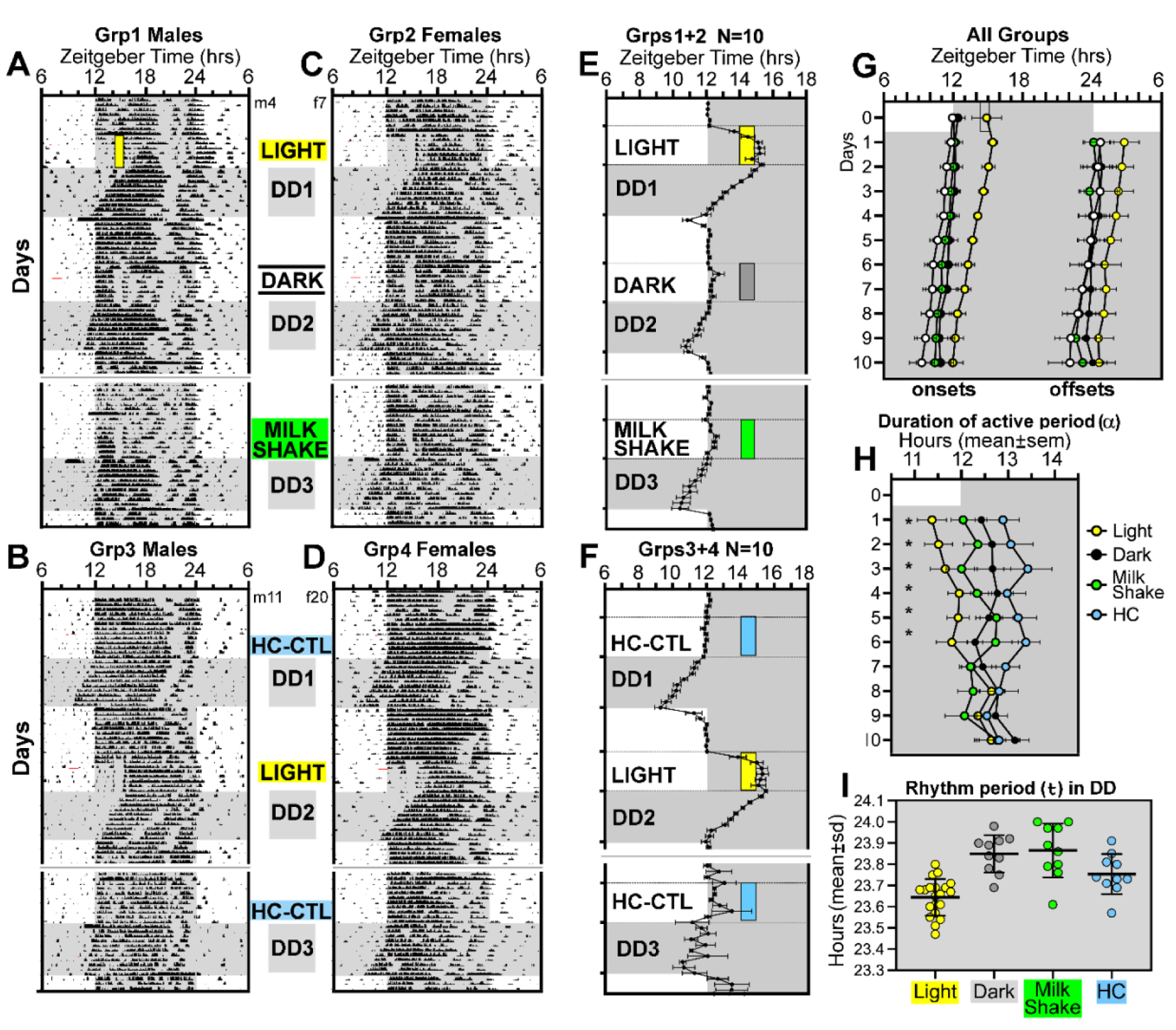
One week of exposure to touchscreen chamber light phase-delays activity onset and shortens free-running period. (**A-D**) Representative actograms of male (**A, B**) and female (**C, D**) mice. (**E, F)** Group mean (**±**SEM) activity onsets for all days in after combining groups 1 and 2, and groups 3 and 4 (**G)** Group mean (**±**SEM) activity onsets and offsets in DD in all four groups. (**H**) Group mean (**±**SEM) activity duration (a, time from onset to offset) in DD in all four groups. \**p* < 0.05. (**I**) Group mean (**±**SD) periodicity (t) of activity rhythms in DD. ZT, zeitgeber time; DD, constant darkness; LD, light:dark.; MS, milkshake; HC, home cage.

Linear regression on the 10 days of activity onsets in DD was used to determine the circadian periodicity (τ) of free-running rhythms. ANOVA revealed a significant main effect of condition on τ (F_3, 24_ = 19.08, *p* < .0001), and no significant effect of sex (F_1, 18_ = 1.304, *p* = .268) or interaction (F_3, 24_ = 0.798, *p* = .507). Tukey’s post-hoc tests indicate that τ was significantly shorter in DD after the Light condition compared to DD after the Dark, Milkshake and Home cage conditions.

The duration of the daily active period (α) in DD also showed a significant main effect of condition (F_3, 24_ = 9.154, *p* = .0003), and no significant effect of sex (F _1,18_ = 1.467, *p* = .2415) or interaction (F_3, 24_ = 1.521, *p* = .2347). Tukey’s post hoc tests indicated a significantly shorter α in the Light condition compared to the Dark, Milkshake and Home cage control conditions. To determine if α was stable in DD, males and females were pooled and a second ANOVA was conducted to evaluate the effect of day. There was a significant main effect of condition (F _3, 46_ = 3.369, *p* = .0264) and day in DD (F_9, 414_ = 1.962, *p* = .0423) and a significant interaction (F_46, 414_ = 10.45, *p* < .0001) on α. Tukey’s post-hoc tests indicate that short α in the Light condition differed from the other conditions only until day 6 in DD. Linear regression on α duration across days in DD yielded a significant positive slope only in the Light condition, indicating that α was compressed by daily chamber light exposure at ZT14.5, and then decompressed in the absence of light exposure. These results indicate that the effect of the Light condition on t derived from activity onsets was likely secondary to decompression of α.

### 3.3 Experiment 3: Lighting modifications to minimize circadian shifting without affecting visual discrimination

Experiment 2 revealed that effects of the TUNL procedure on circadian rhythms in LD were due to touchscreen lighting. Experiment 3 aimed to identify lighting conditions that could minimize these effects while preserving visual discrimination (>70% accuracy). Table 1 describes ten conditions (Standard and A–J) with unique combinations of stimulus luminance (50–100%), color (white or red), screen brightness/backlight (0–50%), and red-green-blue (RGB) channel ratio. Conditions were further characterized by spectrometer measurements at the stimulus location, including illuminance, correlated colour temperature, melanopic irradiance, and chromaticity coordinates (see Supplementary Table S1). Mice underwent touchscreen pretraining as described in Experiment 1 [16, 17], to the ‘Must Initiate’ stage using normal screen conditions. Mice were then tested using a modified trial type referred to as “Correct Touch” in which they had to correctly touch the randomly presented stimulus to get the food reward. Incorrect response resulted in a ‘time-out’ without the house light component. Once mice reached criterion on Correct Touch (with correction trials), mice underwent two days of visual discrimination testing. For each day, mice were tested on three screen conditions (48 trials, 30 min each), with 5-min inter-condition breaks running the Correct Touch schedule under normal screen conditions to prevent extinction. Screen condition order was counterbalanced across mice to control for order effects on performance. Training and testing were conducted at ZT14.5 (2.5 h after lights-off). Visual discrimination performance varied by the screen conditions (one-way ANOVA, F_10,61_ = 13.16, *p* < .0001), with screen conditions A (*q*(61) = 4.430, *p* = 0.0004) and C (*q*(61) = 8.215, *p* < .0001) impairing accuracy relative to baseline performance (Fig. 3A).

**Figure 3.**
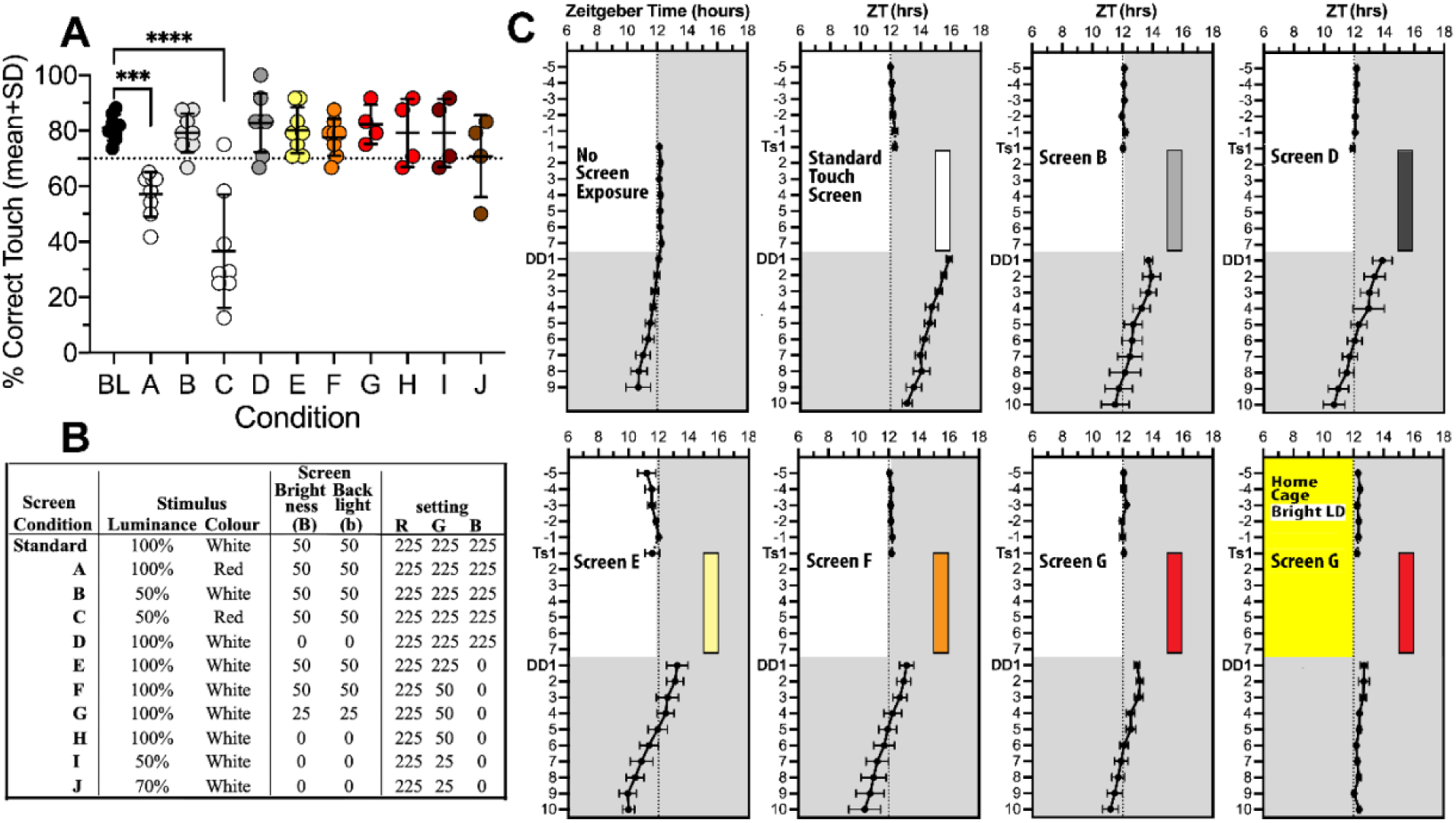
Screen modifications maintain visual discrimination performance while reducing circadian phase shifts. **(A)** Mean accuracy (session % correct) on the Correct Touch visual discrimination task under each of the 10 screen conditions (noted along the x-axis). The 3-day baseline average under standard conditions is highlighted with grey shading. The horizontal dotted line marks the performance criterion (70% accuracy). Each open circle is a single mouse tested under the screen conditions (48 trials, 30 min). Data presented as mean (±SEM). \**p* < 0.05 **(B)** Lighting parameters for each condition. **(C)** Group mean (±SEM) daily activity onsets of mice in 8 screen conditions on pre-exposure baseline days and 10 days of constant darkness (DD). Grey shading denotes lights-off in the home cage. Yellow shading in the screen G panel (lower right) denotes bright home cage lighting. The coloured bars indicate the timing of the daily screen exposures, with bar colour indicating condition to match with panel A.

The mice were then randomly assigned to two groups (Group 1 n=4; Group 2 n=4; females 2/group) and exposed to a modified screen condition for 7 consecutive days (1-h exposures at ZT14.5), followed by DD for 10 days, to assess circadian phase-shifting response. Group 1 was tested with 5 exposure conditions (B, D, F, G, standard) and Group 2 with 4 exposure conditions (E, F, G, and G with bright home cage lighting). Conditions are detailed in Fig. 3B. Mice in the ‘bright home cage lighting’ condition were housed under bright LED light (1000-2000 lx) between ZT0-12 for 7 days before and during the 7-day exposure. There were at least two weeks between exposure conditions to ensure stable LD entrainment between trials.

One week of exposure to the screen conditions caused varying degrees of phase shift (Fig. 3C- J; one-way ANOVA (F_7,36_ = 8.112; *p* < .001). On day 1 of DD, the mean phase delay in activity onset was largest for standard conditions (Fig. 3D, 3.63 h 3.75 **±** 0.43 h). Delays were smaller in conditions B (1.64 **±** 0.54 h), D (1.78 **±** 1.29 h), E (1.80 **±** 0.68 h), and F (0.77 **±** 0.67 h), and smallest in groups F (0.7 **±** 0.67 h), G (1.15 **±** 0.07 h), and G + bright housing light (0.38 **±** 0.43 h).

### 3.4 Experiment 4: TUNL performance under screen conditions that minimize phase shifting

Screen condition G in Experiment 3 substantially reduced phase shifting without impairing visual discrimination and was selected for further testing. Mice were assigned to one of 5 groups (Fig. 4A) with different combinations of touchscreen chamber and home cage lighting intensities. They progressed through pre-training, Cued TUNL, and TUNL as described in Experiment 1 for ∼100 days.

**Figure 4.**
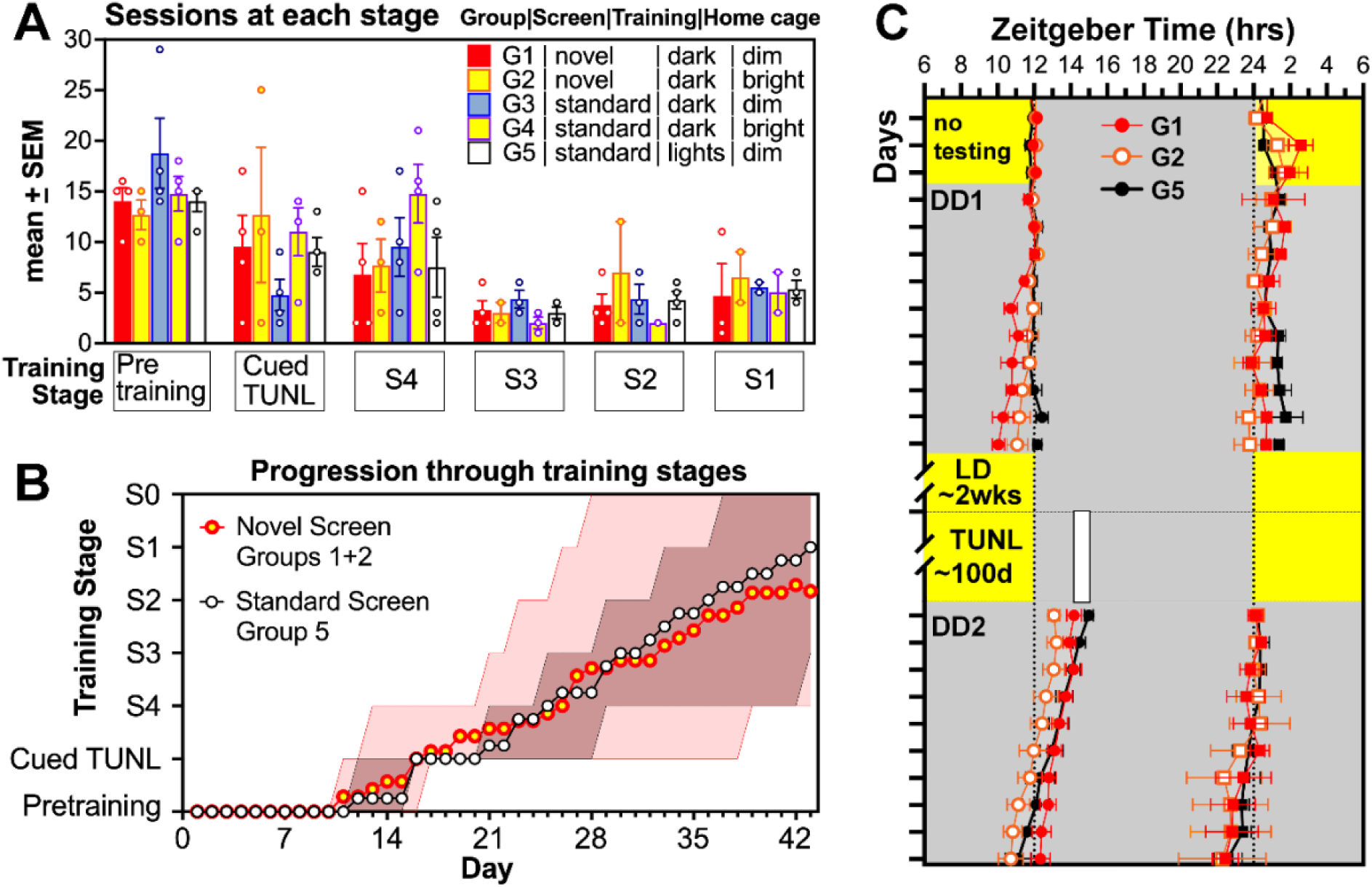
Training on the novel screen (Screen G) does not slow acquisition of TUNL (trial- unique non-matching-to-location). (A) Mean number of sessions to reach criterion performance for pretraining, cued TUNL, and TUNL. The 5 groups differed in touchscreen settings, touchscreen chamber lighting during the task and home cage lighting, as specified in Panel A. Data presented as mean (±SEM); solid circles show data for an individual mouse. (B) Mean training stage reached on each daily session for Group 1 (novel screen G, dark training, dim housing lighting; red) and Group 5 (standard screen, standard lighting in touchscreen chamber, and standard dim home cage lighting; black). The y-axis runs from pretraining through cued TUNL to TUNL levels S4 to S1. Shaded regions show the range of individual mice, pink shading for Group 1 & 2 mice and grey shading for overlap between Group 1 & 2 and Group 5 (only a few mice reached Stage 0, and this is included in the range). (C) Group mean (±SEM) activity onsets (circles) and offsets (squares) for each day of constant dark (DD) following baseline recording in LD and again following ∼100 days of TUNL training in Groups 1 (red), 2 (tangerine) and 5 (black).

Mice trained using screen G (Group 1) or screen G with bright home cage lighting (Group 2) progressed through pretraining, cued TUNL, and TUNL training stages at similar rates to Groups 3–5 that were tested using standard screen conditions (Fig. 4A; mixed-effects analysis showed no significant group difference in sessions completed at each stage F_4, 85_ = 0.1207, *p* = .975). A direct comparison between screen G (Groups 1 and 2 combined) and the standard test conditions (Group 5), indicated similar rates of progression through the training stages to S1 (Fig. 4B). Regression lines fit to each curve confirmed equivalent slopes (F_1, 468_ = 3.516, *p* > .05) and y-intercepts (F_1, 469_ = 0.313, *p* > .5). As expected, phase shifts of activity onset measured in DD were largest in Group 5 (mean ±SD), but were also larger in Groups 1 (mean ±SD), and 2 (mean ±SD), compared to those screen conditions in Experiment 3, likely due to the extended period of training (∼100 days vs 7 days; Fig. 4C).

## 5. Discussion

Touchscreen-based cognitive testing is increasingly used to measure cognition in rodent models [1]; however, to our knowledge the impact of nocturnal testing and the test environment on circadian rhythms have not yet been considered as potential variables. Here we show that daily touchscreen testing can alter the phase of entrainment to the LD cycle and can disrupt re-entrainment to a 6-h LD delay shift. This is a potentially important confound for behavioural testing, for four reasons. First, our results suggest that phase shifting can impair performance on the touchscreen tasks (e.g., TUNL), and thus the unintended phase shifts could be affecting the outcome variables of interest. Additional work with a larger sample size will be necessary to determine whether variability in LD re-entrainment might relate to test performance following an LD shift.

Second, if the circadian clock is inadvertently shifted, and test time remains fixed, then cognitive testing will be occurring at a different circadian phase than intended, e.g., a 3-h delay of the circadian cycle in a mouse tested at ZT14 (2 h after lights off) will effectively move the circadian phase of testing from the third hour of the biological night (daily active phase) to the last hour of the biological day (daily rest phase). The degree of shift may vary within a group, introducing additional uncontrolled variability in circadian test time. Notably, some studies observe variation in learning and memory in mice dependent on circadian timing of training and testing (e.g., [9, 19, 20]).

Third, while exposure to touchscreens early in the dark period produced a clear delay in activity onset, the phase delay was less apparent in the end of activity, indicating compression of nocturnal activity. This is a characteristic of circadian rhythms in nocturnal rodents when the photoperiod (length of the daily light period) is increased. The compression was evident on the first day of DD, after which alpha gradually expanded (or de-compressed). This supports an interpretation that light exposure early in the dark period simulates a long-day photoperiod, which is significant given that photoperiod has been reported to affect performance on behavioural tests under at least some conditions ([21], but see [22]).

Fourth, if the objective is to assess effects of a shifted LD cycle on cognitive processes, then the resistance to shifting evident in some but not other mice, introduces additional uncontrolled (and potentially unknown) variability in the experimental manipulation.

By separating the photic and non-photic components of the testing procedure, we showed that light emitted by the test platform was sufficient to reproduce the phase shifting effect of the full standard TUNL protocol. One week of daily exposure to the touchscreen chamber with standard light settings delayed activity onset by approximately 3.4 h and altered parameters (tau and alpha) of the activity rhythm in DD. Neither the food reward nor chamber exposure in darkness produced these effects, confirming light emitted by the touchscreen chamber as the critical stimulus. Together, these findings show that routine touchscreen testing can alter not only circadian phase but also fundamental properties of the pacemaker, changes that would go undetected in most behavioural studies that do not routinely monitor circadian rhythms.

To determine whether touchscreen lighting could be modified to minimize circadian impact without impairing visual discrimination, we designed 10 unique lighting conditions that varied stimulus brightness and spectral composition, screen backlight, and intensity of light in the home cage. The phase-delaying effect scaled with light intensity and blue wavelength content of the screen condition. The best-performing modifications (Screen G) reduced the delay from ∼3.6 h to <1 h, with a further reduction to ∼0.5 h when combined with bright home cage lighting (bright LED light, 1000-2000 lux). Mice trained under Screen G progressed through pretraining, cued TUNL, and TUNL at rates comparable to mice trained under standard conditions. These results indicate that circadian impacts can be substantially reduced without compromising task acquisition.

Based on these findings, we offer the following recommendations for touchscreen testing of nocturnal rodents during their active phase (Box 1). Relative to the standard configuration, we recommend reducing screen brightness and backlight (from 50 to 25) and applying a 225:50:0 RGB ratio, which removes the blue channel and strongly reduces the green channel. Stimulus luminance can stay at 100%, but the screen output is shifted toward long wavelengths and dimmed while stimulus contrast is maintained. Stimulus colour is kept as “white,” but the stimuli are rendered red-orange by the RGB setting.

Computer-automated cognitive test batteries have been proposed as a way to improve translation from preclinical studies to clinical trials in neurodegenerative and neuropsychiatric disease, largely by increasing the reproducibility and translational validity of cognitive assessment in animal models [23]. Our findings indicate that realizing this potential will require attention to less obvious features of the testing environment, such as the circadian effects of chamber lighting, which can introduce systematic variability even in standardized, automated protocols.

While these findings are relevant to all behavioural testing of mice occurring during the active (dark) phase, they are of particular importance to studies that use touchscreen tasks to examine how sleep and circadian disruption affect cognition. They are also relevant to studies of rodent models of conditions in which circadian dysfunction is a prominent feature, including Alzheimer’s disease and aging [24]. In such studies, the testing procedure could act on the independent variable. Daily light exposure in the chamber could partially re-synchronize a disrupted rhythm, similar to a light-therapy intervention, and attenuate or mask a phenotype. It could also misalign rhythms in otherwise healthy controls, reducing apparent group differences. Similar concerns apply to other operant paradigms that use illuminated stimuli, house lights, or cue lights during the dark phase, and to other nocturnal rodents, such as rats.

Several limitations should be considered. First, because females were pair-housed in Experiment 1 to avoid social isolation stress, individual activity records were not available for those animals. While the aggregate activity rhythms of these pairs were remarkably similar to the rhythms of the single housed animals, suggesting a high degree of social synchrony between paired mice, the aggregate measure could mask shifting of one mouse to the procedure if the other mouse does not shift. Second, we did not directly test whether the modified testing conditions eliminate the impaired re-entrainment response to the 6-h LD shift observed in Experiment 1. We suspect that phase shifting was limited in these cases because the touchscreen procedure had altered the entrained phase of the circadian clock, thereby changing the circadian timing of the LD shift, which is what determines the response of the clock. If this interpretation is correct, then minimizing the phase delaying effect of the touchscreen procedure should normalize the LD re-entrainment response. A third limitation is that although bright home cage lighting further reduced the phase delay produced by Screen G, we did not test whether it alone, without screen modifications, might be a sufficient prophylactic to achieve this objective. Such an approach would be simpler to implement than modifying screen conditions. However, nocturnal rodents are sensitive to light, preferring dim environments, and bright light has been associated with increased anxiety-like behaviour and other behavioural changes [25–27]. The light levels required to produce rhythms insensitive to entrainment by touchscreen light would be difficult to standardize.

### 5.2. Conclusion

Daily touchscreen testing during the active phase can shift circadian rhythms of mice, primarily through light emitted by the chamber. The effect is large enough to alter the phase of entrainment to LD and to impede re-entrainment to a shifted LD cycle. Modifying the chamber by removing short wavelengths, reducing luminance and backlight, and pairing these changes with brighter housing lighting substantially reduced the circadian impact while preserving visual discrimination and TUNL acquisition. These adjustments are particularly important for studies in which sleep, circadian rhythms, or their disruption are central to the research question.

## Acknowledgements

This work was funded by Natural Sciences and Engineering Research Council of Canada Discovery Grant (RGPIN/03909-2021, BAK), Canada Research Chair (CRC-2020-00047, BAK), Canada Foundation for Innovation (CFI-41428, BAK). We would like to thank the SFU animal research staff for their support and care for the animals used in this study.

## Notes

### Competing Interest Statement

The authors have declared no competing interest.

