## Supplementary Table 1 for "Standard touchscreen test procedures shift circadian rhythms in mice: a modified protocol to minimize circadian confounds"

Supplementary Materials

|  | E_v (lx) | E_s (lx) | E_s/E_v (S/P ratio) | CCT (K) | E_e,mel (W/m²) (mel. Irradiance) | Human Ev,mel,D65 (lx) mel EDI Illuminance) | E_e Irradiance (W/m²) | PPFD (μmol/m²s) | x | y | u' | v' | delta uv | Dominant λ (nm) | Peak λ (nm) | Centroid  λ (nm) |
| --- | --- | --- | --- | --- | --- | --- | --- | --- | --- | --- | --- | --- | --- | --- | --- | --- |
| Standard | 8.306 | 16.15 | 1.9441 | 5733.5 | 5.862 | 8.306 | 0.02829 | 0.1155 | 0.327 | 0.3495 | 0.2 | 0.481 | 0.0067 | 528.8 | 881 | 707.6 |
| A | 2.088 | 1.15 | 0.5506 | N/A | 0.3799 | 2.088 | 0.01103 | 0.04276 | 0.5762 | 0.341 | 0.388 | 0.5167 | N/A | 607.1 | 879.6 | 800.8 |
| B | 2.112 | 4.717 | 2.2335 | 7826.7 | 1.8 | 2.112 | 0.009293 | 0.03125 | 0.296 | 0.3095 | 0.1934 | 0.455 | 0.002 | 483 | 879.1 | 795.1 |
| C | 0.7347 | 0.9368 | 1.2752 | 2144.2 | 0.3472 | 0.7347 | 0.005043 | 0.01329 | 0.4524 | 0.3317 | 0.2979 | 0.4914 | 0.0305 | 611.7 | 879.7 | 835.5 |
| D | 1.048 | 2.179 | 2.079 | 6409.4 | 0.8128 | 1.048 | 0.005235 | 0.01523 | 0.314 | 0.3323 | 0.1975 | 0.4703 | 0.0043 | 491 | 879.6 | 825 |
| E | 7.924 | 10.87 | 1.3713 | 3890.9 | 3.183 | 7.924 | 0.02168 | 0.09079 | 0.4248 | 0.5147 | 0.2041 | 0.5563 | 0.0421 | 570.1 | 888.4 | 739.7 |
| F | 2.921 | 2.626 | 0.899 | 1867.1 | 0.8144 | 2.921 | 0.01255 | 0.0495 | 0.529 | 0.3945 | 0.3169 | 0.5318 | 0.0053 | 592 | 879.7 | 788.7 |
| G | 1.534 | 1.265 | 0.8245 | N/A | 0.409 | 1.534 | 0.007703 | 0.02788 | 0.5373 | 0.37 | 0.335 | 0.5231 | N/A | 597.5 | 888.1 | 815.2 |
| H | 0.5654 | 0.5799 | 1.0257 | 1821.4 | 0.2054 | 0.5654 | 0.003848 | 0.01053 | 0.4994 | 0.3503 | 0.3219 | 0.5231 | 0.0214 | 601.9 | 882.6 | 842.5 |
| I | 0.3781 | 0.5057 | 1.3375 | 2252.5 | 0.1887 | 0.3781 | 0.003021 | 0.006875 | 0.4478 | 0.3376 | 0.291 | 0.4936 | 0.0281 | 606.6 | 887.5 | 846.3 |
| J | 0.5186 | 0.4976 | 0.9595 | 1689.1 | 0.1789 | 0.5186 | 0.003797 | 0.009991 | 0.5089 | 0.37 | 0.335 | 0.5053 | 0.0236 | 606 | 879.2 | 843.3 |

Table S1. Spectrometer readings
